# A *tetRA*-based promoter system for the generation of conditional knockouts in *Campylobacter jejuni*

**DOI:** 10.1101/616649

**Authors:** Eli J. Cohen, Rui Tong Quek, Morgan Beeby

## Abstract

*Campylobacter jejuni* is responsible for tens of millions of cases of gastroenteritis each year. Despite its prevalence and impact on human health, the repertoire of genetic tools available for researchers to study *C. jejuni* remains limited. In order to expand upon the genetic toolkit in this species, we have engineered a system for generating conditional knockouts based on the *tetRA* tetracycline-resistance cassette. This system exhibits tight repressibility and titratability of target-gene expression and will be useful for future research on this important human pathogen.

## Introduction

*Campylobacter jejuni*, a gram-negative human pathogen, causes millions of cases of gastroenteritis worldwide each year (1, 2). Additionally, *C. jejuni* is implicated in the development of Guillain-Barre syndrome, a debilitating neurological disorder, in a small subset of patients (3). As such, much research has been performed to understand the molecular mechanisms of pathogenesis by *C. jejuni.*

Despite research on this species going back several decades, the genetic toolkit for *C. jejuni* remains small compared with other important pathogenic species of bacteria. In part, this can be attributed to the difficulty of importing pre-existing tools used in other bacterial species into the ε-proteobacteria, of which *C. jejuni* is a member. For example, promoter structure in *C. jejuni* is sufficiently different from the γ-proteobacteria that functional promoters in *E. coli* are not recognized by *C. jejuni* RNA polymerase (RNAP), and vice versa (4).

In order to expand the portfolio of tools available for use in *C. jejuni*, we set out to develop an inducible-promoter system based on the *tetRA* tetracycline-resistance cassette. The *tetRA* system was originally discovered and characterized as part of the *Tn10* transposon in *Salmonella typhimurium* (5-8) and has since been used as a tool for generating conditional knockouts in several species of bacteria (9-13). The *tetRA* cassette consists of two genes, *tetR* and *tetA*, that are divergently transcribed from overlapping promoters (Fig. 1A). The TetR protein forms a dimer and, in the absence of tetracycline, binds to *tetO* operator sequences in the *tetRA* promoter region, thereby repressing both *tetR* and *tetA*. In the presence of tetracyclines, TetR is released from *tetO* sequences, thereby allowing *tetA*, encoding the tetracycline efflux pump, to be expressed.

**Figure 1:**
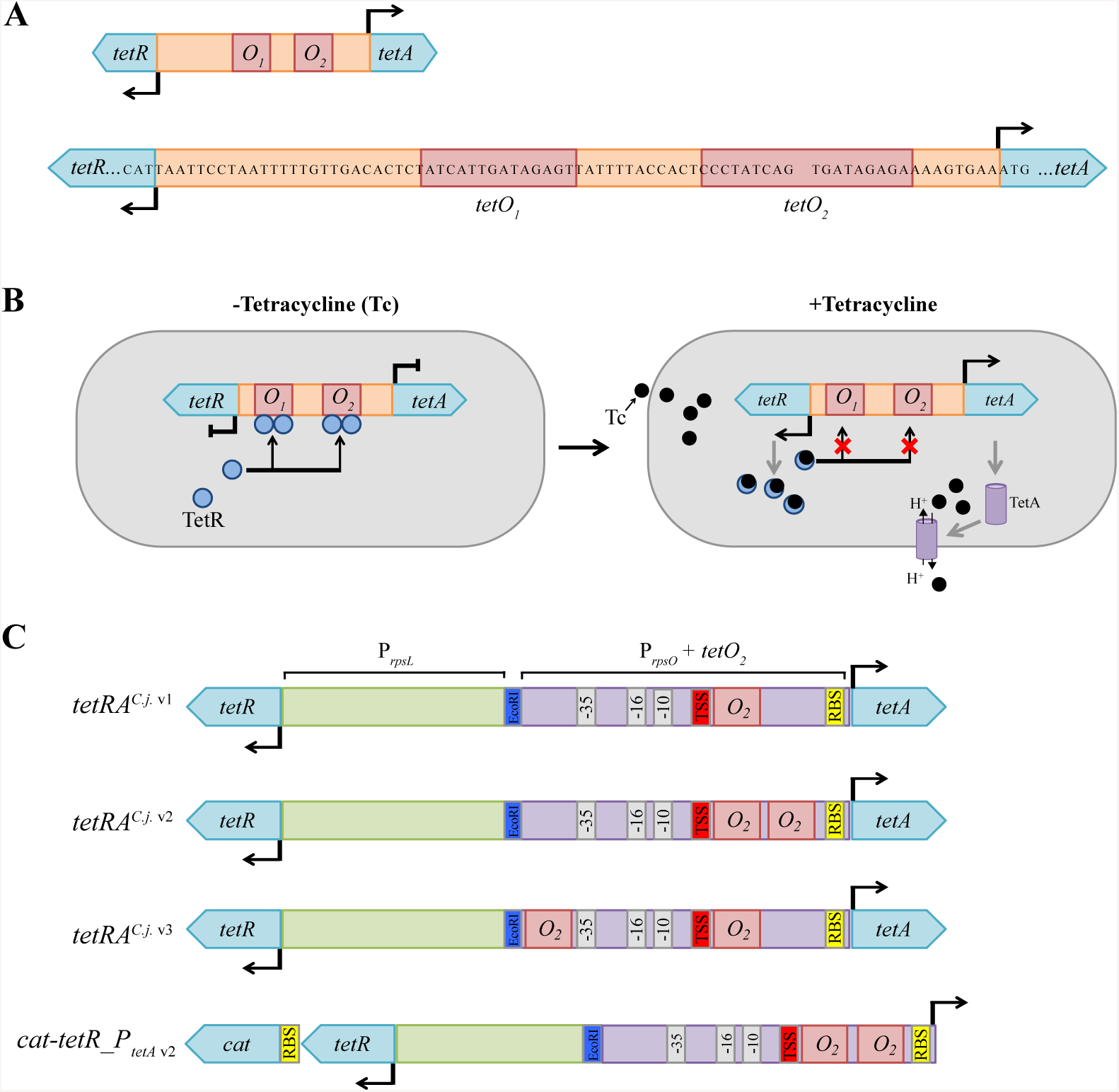
Overview of the Tn10 *tetRA* and architecture of *tetRA*^*C.j.*^ cassettes. The tetRA tetracycline-resistance determinant was first discovered and characterized as part of the Tn10 transposable element in *Salmonella enterica*. **(A)** The Tn10 *tetRA element* consists of the *tetR* gene, encoding the TetR repressor protein, and *tetA*, which codes for the TetA tetracycline-efflux pump. *tetR* and *tetA* are expressed from the 78-bp promoter region containing *tetO*_*1*_ and *tetO*_*2*_ TetR binding sequences. **(B)** In the absence of tetracyline, TetR dimerizes and binds to *tetO* sequences, repressing expression of both *tetR* and *tetA*. Upon addition of tetracycline, TetR is released from the *tetRA* promoter region, thereby allowing the expression of *tetR* and *tetA.* **(C)** The *Campylobacter jejuni tetRA* cassette, *tetRA*^*C.j.* v1^, and its derivatives were constructed by fusing the constitutively-expressed *rpsL* and *rpsO* promoters to one another in divergent orientations. The *rpsO* promoter driving *tetA* was modified by adding *tetO*_*2*_ sequences flanking between the ribosom binding site (RBS) and transcriptional start site (TSS) (v1 and v2), and just upstream of the −35 region (v3). An EcoRI site was added between the two promoters for methylation in order to increase transformation ef ficiency during strain construction. The *cat-tetR_P*_*tetA* v2_ cassettes possess the cat gene transcriptionally fused downstream of *tetR* and have had *tetA* excised. The promoter regions of the *cat-tetR_P* _*tetA*_ cassettes are identical to the *tetRA*^*C.j.* v1-3^ cassettes.

By replacing the native *tetRA* promoters with synthetic, *C. jejuni*-derived promoter regions, we were able to construct a functional *tetRA*-based expression system for use in *C. jejuni*. The *tetRA*-derived cassettes possess tight repressibility as well as titratability of target-gene expression. To the best of our knowledge, this is the first inducible-promoter system developed for use in this species.

## Results

### Design of *tetRA*^*C.j.*^

Using DRH212 (14), a tetracycline-sensitive strain of *C. jejuni* 81-176, as our wild-type (WT) strain, we tested several *tetRA*s harboring engineered promoters for their ability to confer tetracycline resistance when integrated in the chromosome. The synthetic promoters were generated by replacing the entire Tn10 *tetRA* promoter region with *C. jejuni* promoters harboring *tetO*_*2*_ sequences. Of those tested, only one was found to enable growth on media containing tetracycline.

The functional *C. jejuni tetRA, tetRA*^*C.j.* v1^, was generated by placing *tetR* downstream of the *rpsL* promoter and *tetA* under the control of a modified *rpsO* promoter harboring a single *tetO*_*2*_ sequence between its −10 region and ribosome binding site (RBS) (Fig. 1B). *rpsL* and *rpsO* code for protein subunits of the ribosome and are highly expressed in *C. jejuni.* The *rpsL* promoter driving *tetR* transcription does not contain a *tetO*_*2*_ sequence and is therefore constitutively expressed.

### Optimization of *tetRA*^*C.j.*^

In order to test the properties of *tetRA*^*C.j.* v1^, we placed the cassette just upstream of *astA*, encoding arylsulfatase, generating the P_*astA*_::*tetRA* ^*C.j.* v1^ reporter strain. Arylsulfatase catalyzes the release of inorganic sulfate from arylsulfate esters and has previously been used as a reporter for gene expression in *C. jejuni* (15, 16).

*tetRA*^*C.j.* v1^ was found to downregulate *astA* expression in the absence of inducer (Anhydrotetracycline (ATc)), and overexpress *astA* in its presence, relative to WT expression levels (Fig. 2A). However, repression of *astA* by *tetRA* ^*C.j.* v1^ was incomplete, which was evident by cultivating this mutant on media containing a chromogenic arylsulfatase substrate (5-bromo-4-chloro-3-indolylsulfate (XS)) lacking ATc.

**Figure 2:**
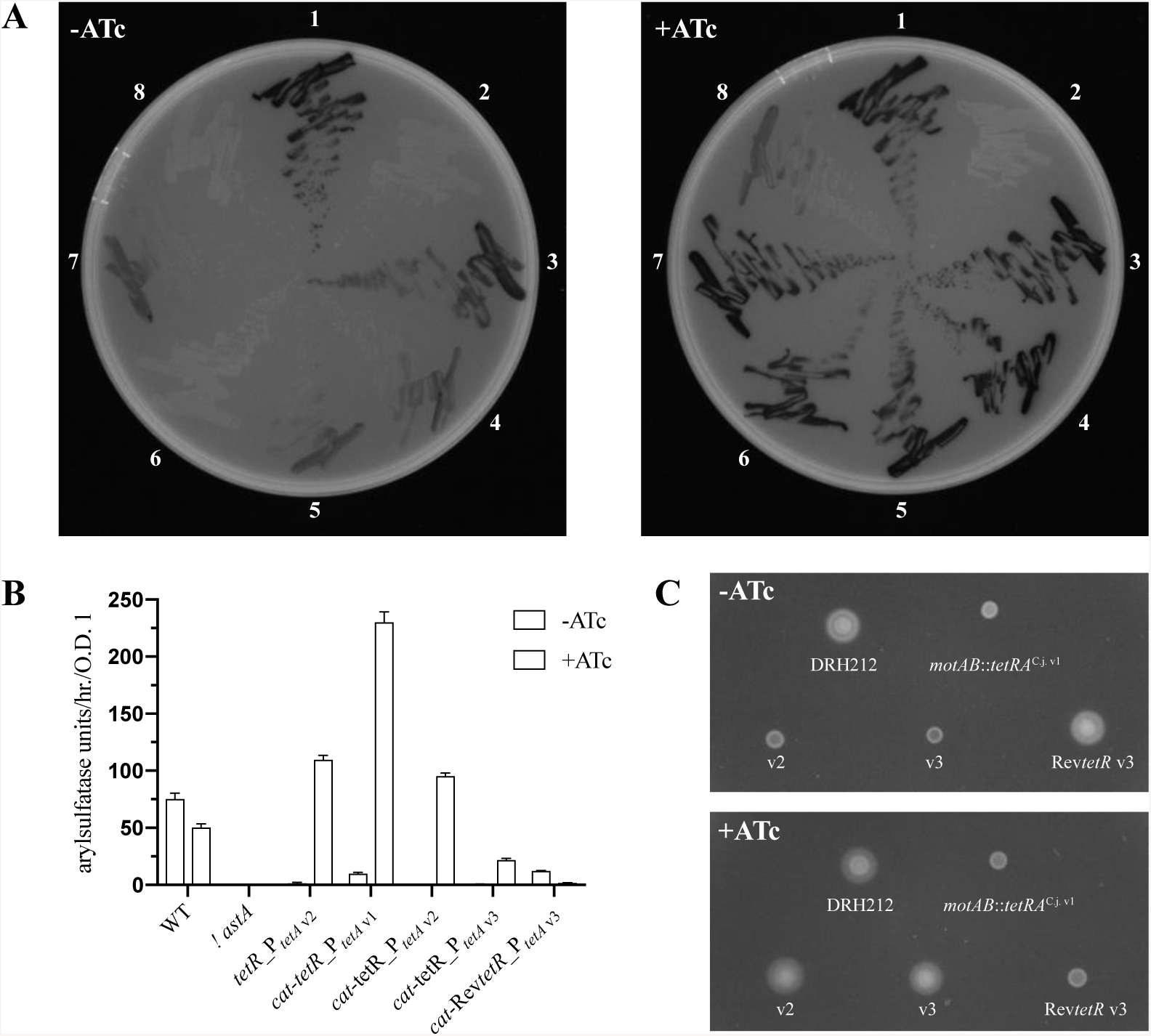
*tetRA*^*C.j.* v1^ and its derivatives regulate target gene expression. The *astA* promoter was replaced with *tetRA*^*C.j.* v1^ and itsderivatives. **(A)** Incubation on XS agar with and without ATc added allowed us to determine whether each cassette was able to regulate *astA* expression and compare the ability of each derivative to repress *astA* expression. Strains: (1) DRH212, (2) □*astA*, (3) *tetRA*^*C.j.* v1^, (4) *cat*-*tetR*_P _*tetA*_ v1, (5) *tetRA*^*C.j.* v2^, (6) *cat*-*tetR*_P _*tetA*_v2, (7) *tetRA*^*C.j.* v3^, (8) *cat*-*tetR*_P_*tetA*_v3. **(B)** AstA activity from the various cassettes with and without ATc added. **(C)** *cat*-*tetR*_P_*tetA*_v2, *cat*-*tetR*_P_*tetA*_v3 and *cat*-Rev*tetR*_P_*tetA*_v3 were tested for their ability to regulate motility by contr olling expression of *flgR*, a response regulator involved in flagellar morphogenesis.

In an attempt to more fully repress *astA* expression in the absence of ATc, two derivatives of *tetRA*^*C.j.* v1^ were constructed: *tetRA*^*C.j.* v2^ and *tetRA*^*C.j.* v3^. Each derivative harbors two *tetO*_*2*_ sequences in the *rpsO* promoter, in different locations (Fig. 1B). Insertion of a second *tetO*_*2*_ site resulted in tighter repression in both P_*astA*_::*tetRA* ^*C.j.* v2^ and P_*astA*_::*tetRA* ^*C.j.* v3^ relative to P_*astA*_::*tetRA* ^*C.j.* v1^ on XS agar without ATc. P_*astA*_::*tetRA* ^*C.j.* v2^ demonstrated somewhat higher *astA* expression than P_*astA*_::*tetRA* ^*C.j.* v3^ on XS agar with ATc added. However, both derivatives still exhibited identical, low-level *astA* expression phenotypes in the absence of ATc (Fig. 2A).

The observation that P_*astA*_::*tetRA* ^*C.j.* v2^ and P_*astA*_::*tetRA* ^*C.j.* v3^ exhibited different phenotypes on XS agar containing ATc, but the same appearance on XS agar lacking ATc suggested that an undesirable level of gene expression in the uninduced state was intrinsic to the *tetRA*^*C.j.*^ cassettes. We presumed that this was due to either incomplete repression of the synthetic *rpsO* promoters by TetR or the presence of cryptic promoter elements within the *tetA* coding region that were being recognized by *C. jejuni* RNAP.

To determine if the leakiness of *tetRA*^*C.j.*^ cassettes could be attributed to background readthrough from within the *tetA* sequence, *tetA* was deleted from all three *tetRA*^*C.j.*^ variants. We found that by deleting *tetA*, we were able to reduce *astA* expression below detectable levels on X-S agar in both the P_*astA*_::*tetRA* ^*C.j.* v2^ and P_*astA*_::*tetRA* ^*C.j.* v3^ backgrounds (Fig. S2).

However, excision of *tetA* sacrificed an advantageous feature of the *tetRA*^*C.j.*^ cassettes: the ability to generate conditional-knockout mutants in the chromosome in one step by selecting for tetracycline resistance. This was addressed by generating a transcriptional fusion of the gene encoding chloramphenicol acetyltransferase, *cat*, to *tetR*. Thus, insertion of the derived cassettes, *cat*-*tetR*_P_*tetA*_^v1-3^, can be selected for on media containing chloramphenicol. Comparison of *astA* expression in P_*astA*_::*tetR_*P_*tetA*_^v2^ and P_*astA*_::*cat*-*tetR*_P_*tetA*_^v2^ genetic backgrounds showed that addition of the *cat* gene did not disrupt the function of the cassettes (Fig 2B).

We next constructed a conditional knockout of *flgR* in order to confirm the results obtained at the *astA* locus. The FlgR protein is a response regulator that links completion of the *C. jejuni* flagellar type three secretion system (fT3SS) to the subsequent expression of genes required for assembly of the flagellar hook basal body (16). Mutants lacking a functional copy of *flgR* are non-motile due to their inability to construct a flagellum.

*cat*-*tetR*_P_*tetA*_^v2^ and *cat*-*tetR*_P_*tetA*_^v3^ were inserted in the chromosome directly upstream of the *flgR* start codon and tested for their ability repress motility in the absence of ATc. Motility agar with and without ATc added were inoculated with the conditional-knockout mutants and allowed to incubate for 12 hours. We found that both *cat*-*tetR*_P_*tetA*_^v2^ and *cat*-*tetR*_P_*tetA*_^v3^ repressed motility in media lacking ATc, and produced motility swarms of approximately WT diameter in the presence of ATc.

In addition to the *cat*-*tetR*_P_*tetA*_^v2^ and *cat*-*tetR*_P_*tetA*_^v3^ derivatives of *tetRA*^*C.j.* v1^, we also constructed a cassette with the *tetR*^V99E^ mutation with the v3 promoter, *cat*-Rev*tetR*_P_*tetA*_^v3^. The V99E mutation converts TetR to a co-repressor in the presence of ATc, i.e. TetR^V99E^ binds to *tetO* sequences in the presence of ATc and represses downstream gene expression. When inserted upstream of *flgR*, the *cat*-Rev*tetR*_P_*tetA*_^v3^ repressed motility in the presence of ATc (Fig. 2C)

### Gene expression from *cat*-*tetR*_P_*tetA*_ is titratable

An attractive feature of *tetRA*-derived promoter systems is the ability to upregulate and downregulate expression of the target gene. To determine if gene expression was titratable using the *cat*-*tetR*_P_*tetA*_ cassettes, we incubated the P_*astA*_::*cat*-*tetR*_P_*tetA*_^v2^ mutant on media containing ATc in concentrations ranging from 0 to 0.4 μg/mL.

Following overnight incubation on varying concentrations of ATc, AstA activity was measured as a proxy for *astA* expression. We found that AstA activity ranged from undetectable levels when incubated without ATc to a peak of ∼70 arylsulfatase units/hr. at a concentration of 0.3-0.4 μg/mL, similar to the WT expression level in DRH212 (Fig. 3B). Cultures grown on media containing lower concentrations of ATc exhibited correspondingly lower AstA activity, indicating that expression of genes under control of the *cat*-*tetR*_P_*tetA*_ cassettes is titratable. Similar results were obtained by western blot probing for FlgR protein in the P_*flgR*_::*cat*-*tetR*_P_*tetA*_^v2^ and P_*flgR*_::*cat*-*tetR*_P_*tetA*_^v3^ backgrounds grown on varying concentrations of ATc (Fig. 3A).

**Figure 3:**
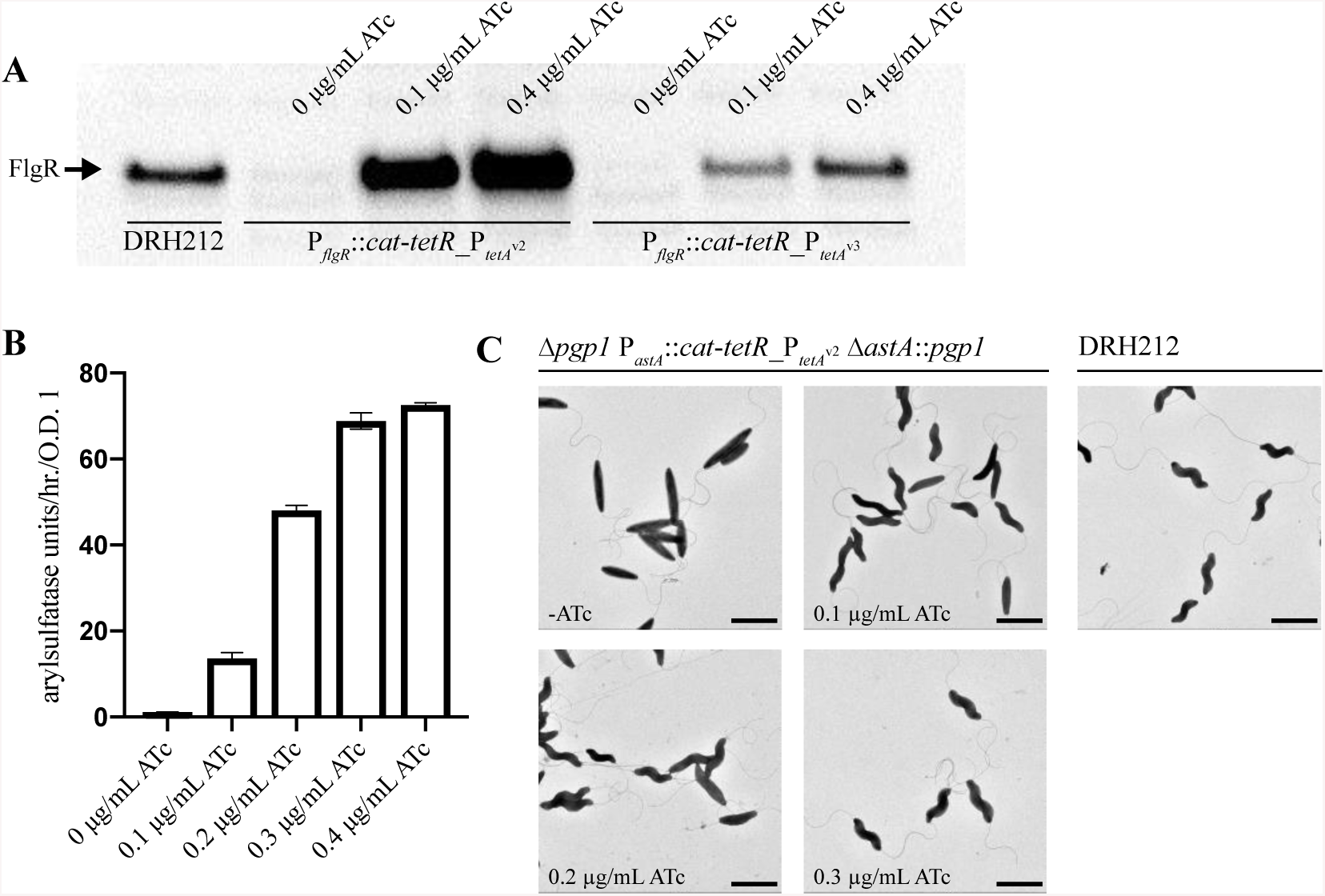
Gene expression from the *cat-tetR_P*_*tetA*_ cassettes is titratable. Three genes, *flgR, astA* and *pgp1*, were used to determine if gene expressi on from the *cat-tetR_P*_*tetA*_ promoter. **(A)** Expression of flgR was tunable using both the *cat*-*tetR*_P_*tetA*^v2^_ and *cat*-*tetR*_P_*tetA*^v3^_ cassette. In the absence of ATc, FlgR was not detected by western blot. In creasing the amount of Atc led to an increase in FlgR expression for both promoters. **(B)** AstA activity was nearly undetectable in the P_*astA*_::*cat-tetR*_P_*tetA*_v2 background in the absence of ATc, and was found to peak at a concentration ∼0.3 μg/mL ATc. Error bars represent the average +/- SEM from three re plicates**(C)** By placing *pgp1* under the control of the *cat-tetR*_P_*tetA*_v2 cassette, the cell body morphology was found to range from a Δ*pgp1* straight-body phenotype in the absence of ATc to a WT helical phenotype at 0.3 μg/mL ATc. At lower levels of ATc, a mixture of cell-body morphologies was observe d. Scale bar = 2 µm.

To confirm the results obtained with *astA*, we decided to test the functionality of the cassette on another gene, *pgp1. C. jejuni* owes its helical cell shape to the action of a handful of enzymes, including Pgp1, that determine peptidoglycan architecture (17). Loss of Pgp1 activity results in a straight, non-helical body morphology.

We generated a markerless deletion of *pgp1* that leaves the first and last 52 codons of the gene intact in order to reduce the likelihood of polarity on neighboring genes. As expected, the Δ*pgp1* mutant exhibited a straight-body morphology but otherwise appeared normal (i.e. WT growth rate, flagella number and cell size). Into this background we inserted a copy of *pgp1* under the control of the *cat*-*tetR*_P_*tetA*_^v2^ cassette at the *astA* locus, resulting in the Δ*pgp1* P_*astA*_::*cat*-*tetR*_P_*tetA*_^v2^ Δ*astA*::*pgp1* mutant.

The *pgp1* conditional knockout strain was incubated overnight on media containing a range of ATc concentrations, from 0 to 0.3 μg/mL, and subsequently observed by transmission electron microscopy. Cells grown in the absence of ATc exhibited a Δ*pgp1* straight-body phenotype, while those incubated on ATc displayed helical body morphologies (Fig. 3C). At the lowest level of ATc tested (0.1 μg/mL), we observed cell shape phenotypes that were neither WT nor that of a *pgp1* knockout, but intermediate between the two. Cells grown on 0.3 μg/mL ATc had a WT cell shape.

### The *cat*-*tetR*_P_*tetA*_ cassette is functional in multiple strains of *C. jejuni*

*C. jejuni* 81-176, a highly invasive clinical isolate, is one of several strains commonly used for research purposes. To test whether the *cat*-*tetR*_P_*tetA*_ cassettes are functional in other strains of *C. jejuni*, we constructed a plasmid, pEJC1, harboring the *cat*-*tetR*_P_*tetA*_^v2^ cassette driving GFP expression. pEJC1 was conjugated into two strains of *C. jejuni*, PT14 and 11168H, and tested for GFP expression in the presence and absence of ATc. When incubated on media lacking ATc, we did not observe fluorescence in either background (Fig. 4). In the presence of ATc, both strains fluoresced, suggesting that the *cat*-*tetR*_P_*tetA*_ cassettes can be used in all strains of *C. jejuni*.

**Figure 4:**
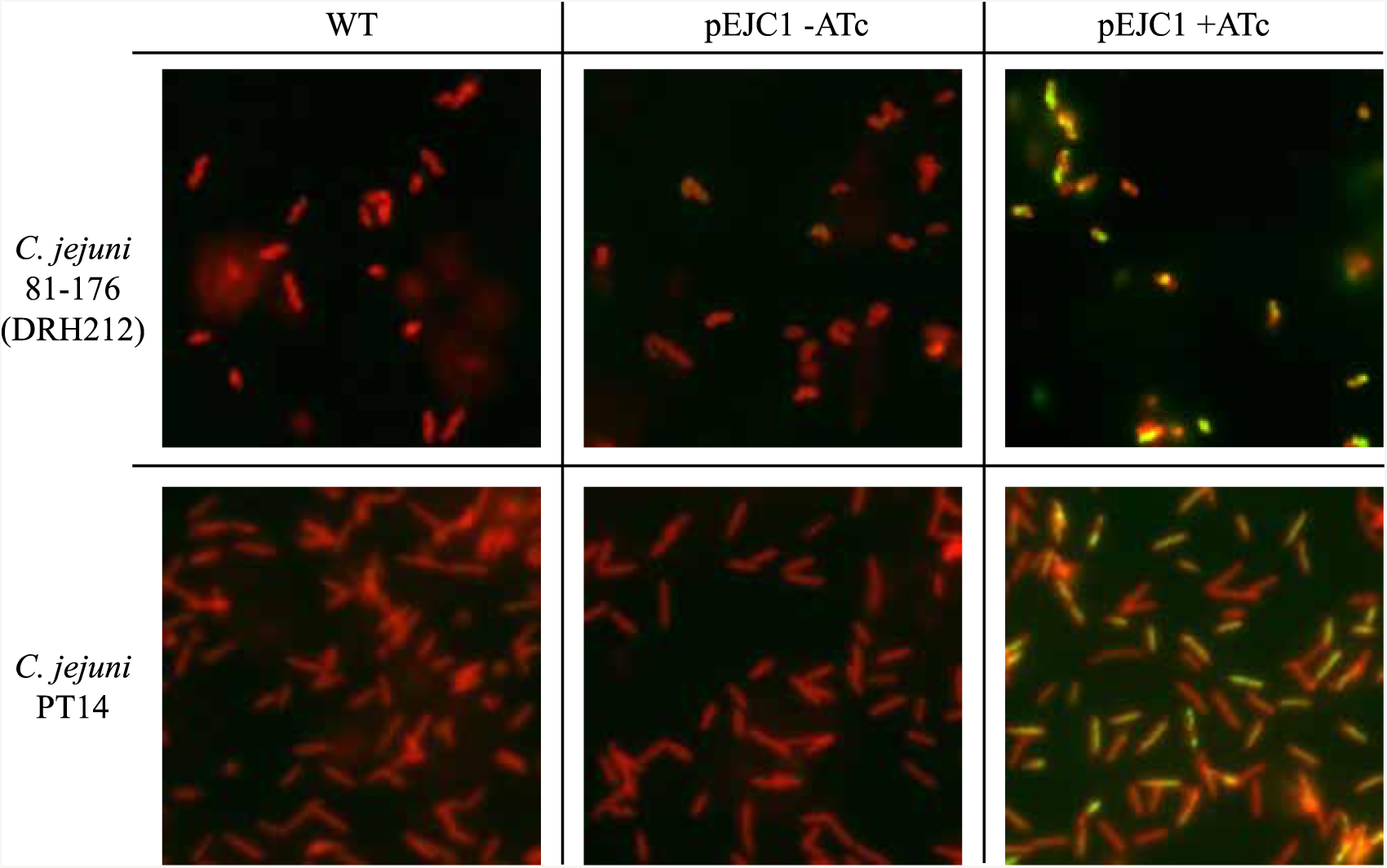
*tetRA*^*c.j.*^ is functional in different strains of C. jejuni. The *cat-tetR_P*_*tetA*^v2^_ driving GFP expression was cloned into pRY108, yielding pEJC1,and conjugated into both DRH212 and C. *jejuni* strain PT14. In the absence of Atc, GFP expression was repressed in both backgrounds. When grown in the presence of Atc (0.4 μg/mL), both strains exhibited fluorescence.

## Discussion

A multifaceted genetic toolkit is required in order to dissect the molecular mechanisms that underpin the behavior and physiology of any organism. Through the efforts of researchers over several decades, a number of genetic tools have been developed that allow for the genetic manipulation of *C. jejuni*. In the work presented here, we have added to that toolkit the ability to generate conditional knockouts and to titrate gene expression in this important human pathogen.

The *tetRA* system was chosen for several reasons. First, *tetRA* gene-expression systems exhibit tight repressibility in the absence of inducer as well as titratability of target-gene expression. Furthermore, the inducers used (e.g. ATc) are membrane permeable, obviating the need for importers of the inducing molecule (e.g. LacY for IPTG). Not only does this reduce the size and complexity of *tetRA* systems, but also allows for target-gene induction across multiple cell membranes, a useful feature when studying intracellular pathogens (such as *C. jejuni*) (9, 11).

Although the strain of *C. jejuni* we used for this work, DRH212, is sensitive to tetracycline, many strains of *C. jejuni* display tetracycline resistance. Resistance in these strains is conferred by the *tetO* gene (different from the *tetO* operator sequences found in the *tetRA* promoter region) either carried on a plasmid or encoded in the genome (18). The TetO protein confers resistance by preventing tetracycline from binding to the ribosome, as opposed to TetA, which is a tetracycline-efflux pump. An advantage of the *tetRA*^*C.j.*^-derivative cassettes, *cat-tetR_P*_*tetA*_^v1-3^, is the ability to generate conditional knockout mutants in both tetracycline-sensitive and resistant strains of *C. jejuni* in one step by selecting for chloramphenicol resistance.

The *tetRA*^*C.j.* v1^-derivatives, *cat*-*tetR*_P_*tetA*_^v2^ and *cat*-*tetR*_P_*tetA*_^v3^, display tight repressibility, titratability and gene-expression profiles over physiologically relevant ranges for the genes studied here, and are currently being used for several projects in our lab. However, we anticipate that it would be relatively straightforward for other researchers to construct *tetRA*^*C.j.*^-based systems with customized expression profiles depending on their needs. We tested only three native *C. jejuni* promoters in our initial attempt to generate a functional *tetRA*^*C.j.*^ cassette, suggesting that this approach would work with many other promoter sequences.

## Materials and Methods

### Bacterial strains and growth conditions

Strains of *C. jejuni* were routinely grown microaerobically (5% O_2_, 10% CO_2_, 85% N_2_) at 37° C on Mueller-Hinton (MH) agar supplemented with 10 μg/mL of trimethoprim. When necessary, media was supplemented with additional antibiotics at the following concentrations: kanamycin: 50 μg/mL, chloramphenicol: 7.5 μg/mL, streptomycin: 2 mg/mL, tetracycline: 5-10 μg/mL (concentration varied by mutant), anhydrotetracycline (0.05-0.4 μg/mL)

### Genetic manipulation of *C. jejuni*

Chromosomal mutations in *C. jejuni* were constructed by transforming linear pieces of DNA generated by splicing-by-overhang-extension PCR (SOE PCR). Individual fragments were designed that possessed 5’ and/or 3’ sequence homology to one another using Q5 high-fidelity DNA polymerase (New England Biolabs), followed by stitching the individual fragments together in a SOE reaction.

Following synthesis of DNA, fragments were transformed into *C. jejuni* using the biphasic method. Briefly, a 5 mm plate with 5 mL MH agar was overlaid with 5-10 μg DNA. Then, *C. jejuni* grown overnight on MH agar was suspended to an O.D._600_ of ∼0.5 in MH broth. 1 mL of cell suspension was added to the DNA-seeded plate and allowed to incubate microaerobically at 37° C for 6-8 hours before being transferred to appropriate selective media and allowed to incubate 3-5 days.

For construction of the markerless Δ*pgp1* strain, an *aph3’-rpsL* (kan-*rpsL*) cassette was first inserted into *pgp1* in DRH212. Insertion of kan-*rpsL* into *pgp1* was selected for via kanamycin resistance. Next, a linear fragment generated by SOE encoding a clean deletion of ∼350 codons of *pgp1* flanked by ∼500 bp of upstream and downstream homology to the desired deletion was transformed into the Δ*pgp1*:: kan-*rpsL* intermediate strain, followed by incubation on MH agar supplemented with 2 mg/mL streptomycin for four days. Streptomycin-resistant colonies were screened for kanamycin sensitivity and sequenced to ensure proper construction of the *pgp1* clean deletion.

### Arylsulfatase activity assays

For determination of *astA* activity on XS agar, MH agar was supplemented with 100 μg/mL XS (5-bromo-4-chloro-3-indolylsulfate (Sigma-Aldrich)) with or without 0.35 μg/mL anhydrotetracycline (ATc) added. Strains to be tested were grown on MH agar from frozen glycerol stocks for 36-48 hours before being struck out on XS agar, followed by an additional 72 hours of incubation.

For spectrophotometric determination of *astA* activity, strains were grown from frozen glycerol stocks for 36-48 hours before being transferred to MH agar with varying amounts of ATc added, followed by an additional 16-20 hours incubation. The strains were then measured for arylsulfatase activity as described previously. Briefly, cells were washed in PBS and resuspended in arylsulfatase buffer (100 mM tris, 1 mM tyramine hydrochloride, 10 mM nitrophenylsulfate, pH 7.2) and incubated for 1 hour at 37° C. Blank measurements were made using cell suspensions incubated 1 hour at 37° C in arylsulfatase buffer lacking tyramine (100 mM tris, 10 mM nitrophenylsulfate, pH 7.2). After one hour, reactions were stopped by addition of stop solution (0.2 M NaOH) before measurement of absorption at 410 nm. 410 nm absorption readings were converted to enzymatic activity by comparison to a nitrophenol standard curve. Arylsulfatase activity values presented in Fig. 2 and 3 are the average of 3 measurements +/- SEM.

### Motility assays

Comparison of motility between DRH212 (WT) and the P_*flgR*_::*cat-tetR_*P_*tetA*_^v2^ and P_*flgR*_::*cat-tetR_*P_*tetA*_^v3^ strains was made using 0.4% agar MH plates with and without ATc added (0.2 μg/mL). Strains were grown from freezer stocks as before and suspended to an O.D._600_ of 1.5. 1μL of cell suspension was spotted on the motility agar and allowed to incubate for ∼12 hours.

### Western Blotting

For detection of FlgR in the *P*_*flgR*_::*cat*-*tetR*_P_*tetA*_^v2^ and *P*_*flgR*_::*cat*-*tetR*_P_*tetA*_^v3^ backgrounds, cultures were grown from freezer stocks for 48 hours on MH agar followed by inoculation to MH agar with varying concentrations of ATc added (0-0.4 μg/mL) and incubated overnight. following overnight incubation, cells were harvested from the plates, washed 1x in PBS and resuspended to an O.D._600_ of 0.8. These were then pelleted and resuspended in 200 μL 2x SDS-PAGE loading buffer. 5 μL of each sample were loaded to each lane of a 4-12% bis-tris gel. Blots were probed for FlgR using FlgR antisera raised in rabbit and developed using HRP-conjugated goat-antirabbit secondary antibody with a ChemidocMP imager (BioRad)

### Electron microscopy

Cells were harvested from MH agar with or without ATc added and suspended to an O.D._600_ ∼1.0 in PBS. 1 mL of the cell suspension was pelleted and resuspended in 1 mL of fixing buffer (100 mM cacodylic acid, 2% glutaraldehyde, pH 7.4). Fixed samples were applied to glow-discharged, carbon-coated copper grids (Agar scientific) and stained with 2% Uranyl Acetate. Micrographs were recorded on a FEI Tecnai T-12 Spirit 120 KeV electron microscope at 4400x magnification.

### Fluorescence microscopy

For observation of fluorescence in strains harboring the pEJC1 plasmid, cultures were grown overnight on MH agar supplemented with 7.5 μg/mL chloramphenicol with or without ATc (0.4 μg/mL) added. Cells were harvested from the plates in PBS and stained with FM 4-64 membrane stain, followed by a wash step then resuspended to an O.D._600_ of ∼1.0 in PBS. Cells were applied to a poly-lysine coated slide and observed on a Leica DMi8 widefield microscope.

## Acknowledgments

We would like to thank Dr. David Hendrixson, Dr. Deborah Ribardo and other members of the Hendrixson lab at UT Southwestern for helpful discussions and advice as well as strains, plasmids and FlgR antisera.

We also wish to thank Dr. Ian Connerton at the University of Nottingham for providing us with strain PT14 and Paul Simpson for technical assistance with the electron microscopy performed for this work.

## List of strains used in this study

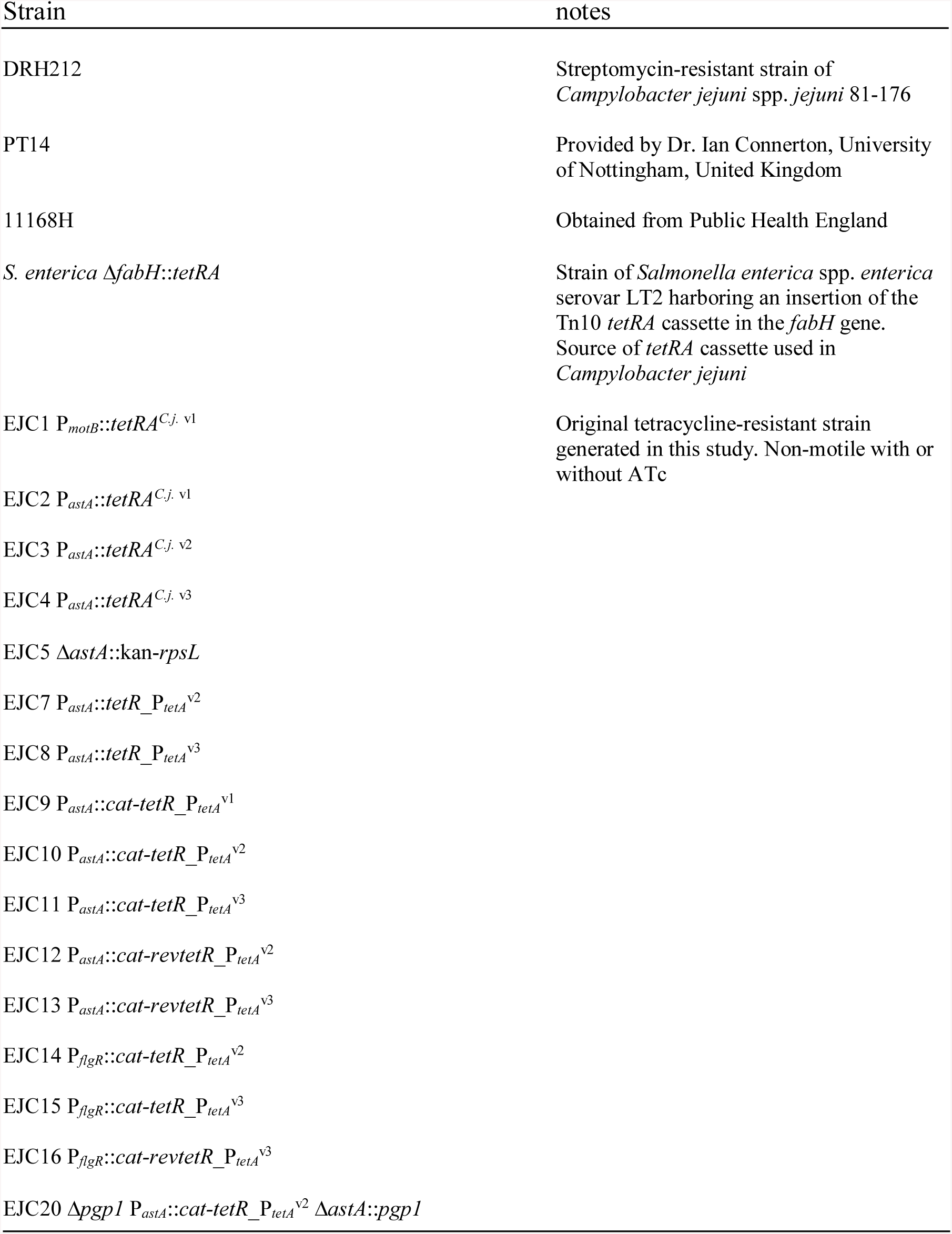

## Primers used in this study

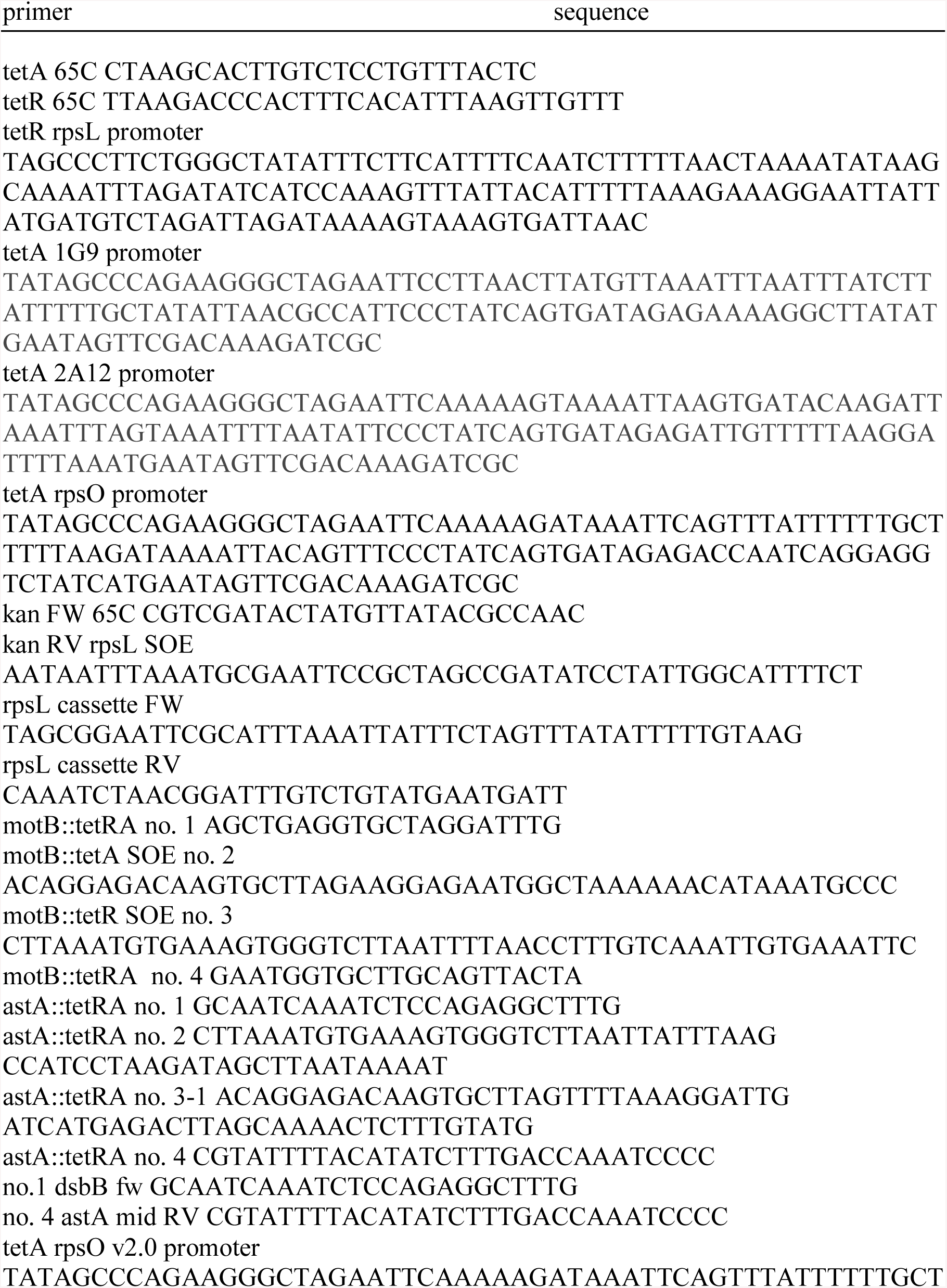
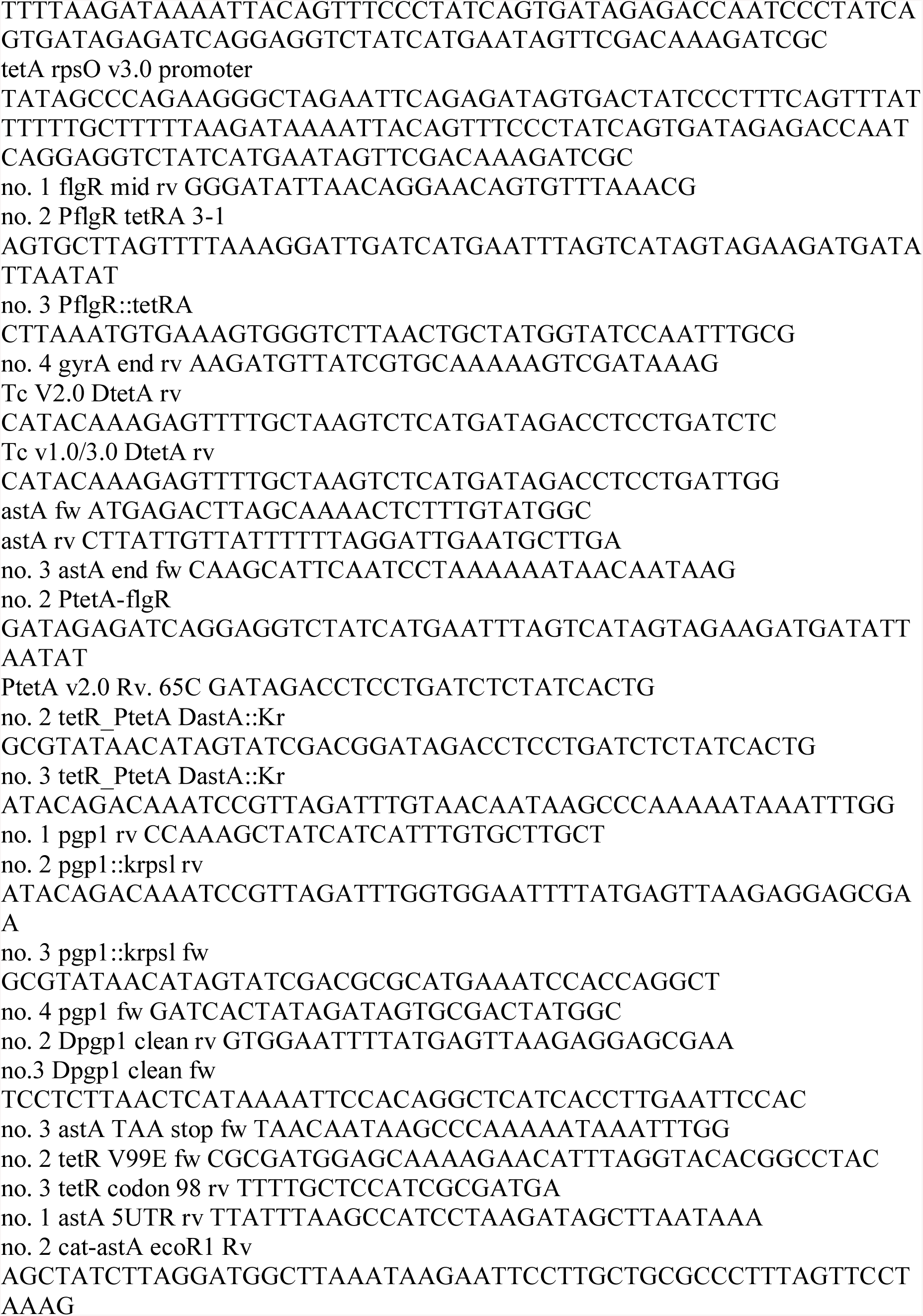
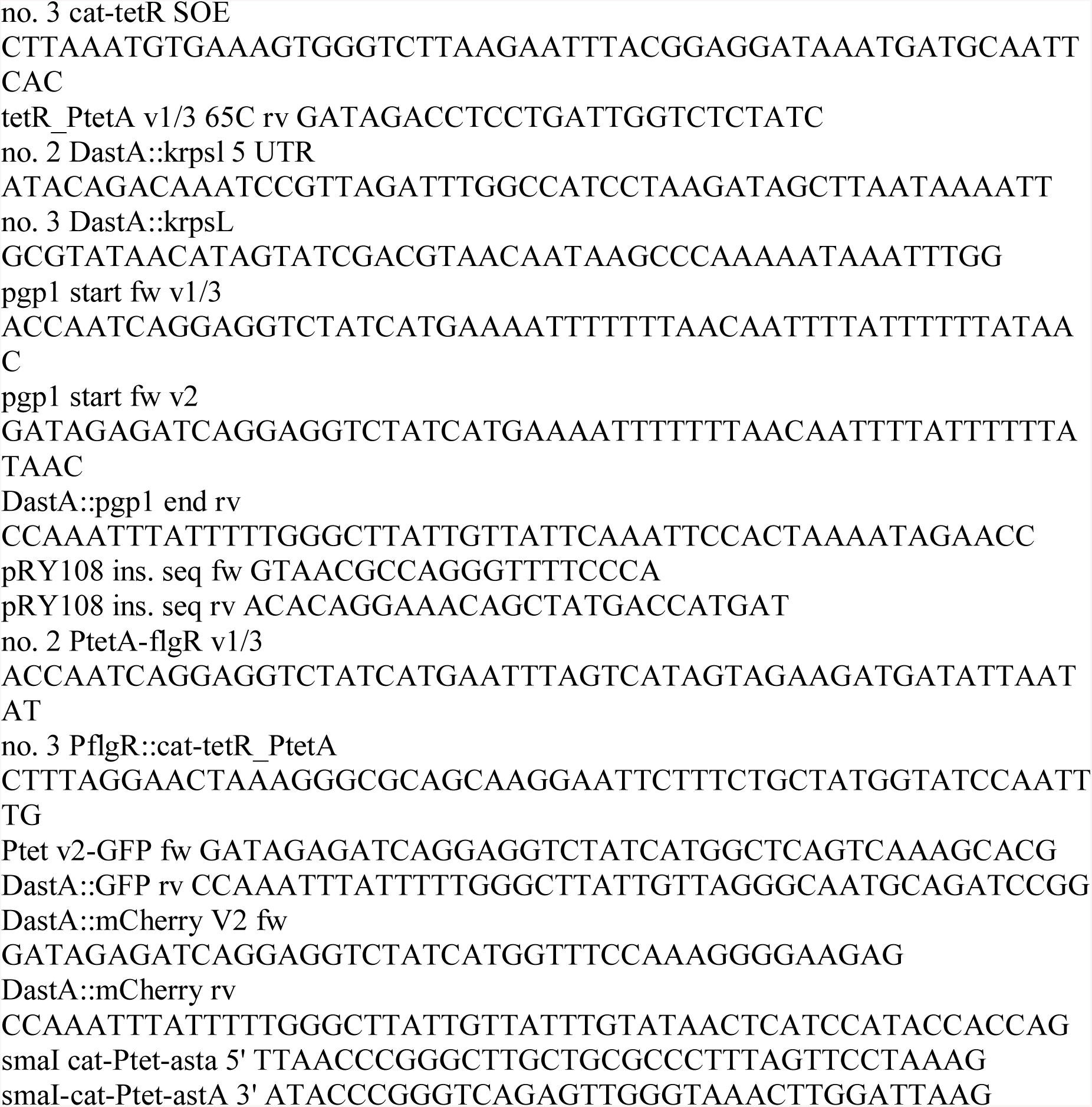

